# Genome-wide analysis of cell-gene interactions

**DOI:** 10.1101/113001

**Authors:** S. Cardinale

## Abstract

The study presents an analysis of how different cellular functions link cell size to the expression of synthetic genes in *E. coli*. The Size-Expression interaction was mapped with a two-gene genetic probe across 3800 single-gene deletion strains. Through regression analysis, expression-specific effects and gene-specific effects were derived from size effects and generic expression effects, respectively. The entire compendium of cell functions broadly mapped to four systems of distinct primary influence on the Size-Expression map. Specifically, membrane structural components primarily affected size, whereas protein and RNA stability primarily affected gene expression. In addition, major Size-Expression shifts showed no substantial gene-specific effects unless they were mediated by key components of the protein synthesis apparatus.

**Subject Category:** Synthetic Biology

## Background

The goal of synthetic biology to standardize the engineering of biology while scaling the design of novel cell functions to increasing complexities critically relies on the reliability and predictability of synthetic biological components ^1,2^. Challenged by constantly mutating endogenous interactions, designed synthetic molecules with specific functions can be quickly made unstable or non-functional by variations in host physiology or genetic makeup. For these reasons, tools are being constructed to shield the functions or predict the behaviors of synthetic genes in the cell. These tools include the design of robustness to evolutionary instability ^3^ and molecular context ^4^ in addition to the application of computational algorithms in the design of molecular parts to ensure robust function ^5^.

Genome-wide mapping of the gene-to-phenotype relationship has enabled the effective identification of genetic targets to improve complex traits in bacteria, such as tolerance to ethanol ^6^, cellulosic hydrolysate and isobutanol ^7^. However, knowledge of the genotype-phenotype relationship does not necessarily allow engineering of reliable heterologous pathways in these strains. To achieve this capability, a systematic understanding of cell-function-to-pathway interactions enabling accurate whole-cell models of the biochemical networks is necessary ^8^.

Here, the cell function-to-gene expression interaction was investigated for all non-essential genes in the *E. coli* genome. Specifically, both the global and specific effects of the lack of a cellular component on cell biomass and synthetic gene expression were mapped. The global effects may involve both the ability of the cell to grow and the cellular amount of a synthetic component. Perturbations in the synthetic circuit dynamics of this type of effect may include growth feedback and have been described mathematically ^9^. Alternatively, a cell function can have a localized effect, such as an effect on the output of synthetic genes but not on cell size. Therefore, to investigate the sequence-specificity of the effects of cell function loss, the present study used a genetic construct with two identically expressed reporter genes.

The results indicated that most functional disruptions affected both cell size and synthetic gene expression and caused an analogous positive or negative shift. Phenotypic patterns of the cell-expression interaction were mapped to 4 major systems in the cell: membrane function, RNA-ribosome-protein stability, biosynthetic and energy-motility functions. Although both sides of the interaction were affected in general, the results suggested that each system predominantly affected one side and indirectly affected the other. In particular, the impairment of cell invagination caused larger cells and indirectly resulted in higher cellular reporter concentrations, whereas defective motility and nutrient uptake yielded mainly smaller cells and secondarily decreased gene expression. Protein folding and RNA stability mainly affected synthetic gene expression while indirectly affecting cell size, particularly if the disruption involved structural ribosome components.

## Results

### Quantification of cell size and reporter expression across the KEIO collection

The aim of this study was to map cell-expression interactions by quantifying the effects of systematic removal of each non-essential *E. coli* gene on the constitutive expression of two synthetic reporter genes. To achieve this, a genetic probe containing the mVenus and mCherry genes expressed from identical promoter-5’-UTR sequences ^10^ was introduced in 3823 gene knockout strains of the KEIO collection ^11^. Each of the 3823 strains carrying the probe was grown from a mixture of 2-3 single colonies from agar plates to avoid colony-to-colony variability of growth on agar and inoculation. Single-cell measurements of mVenus and mCherry fluorescence along with other parameters were acquired with a flow cytometer at the mid-log phase of growth. The variability across biological replicates and data collected at different times was tested for 180 strains from three different plates and was found to be 5*10^−4^ for mVenus and 1.3*10^−3^ for mCherry. These values were approximately 1 order of magnitude smaller than the variance across KEIO strains for the measured variables, which was between 3.6*10^−3^ and 1*10^−2^ (Sup. Info). Although forward scattered light (abbreviated FSC) can be effectively used to measure microbial cell size in cytometry ^12^, this measure does not scale linearly with particle sizes for all instruments ^13,14^. In this study, the cytometer FSC measure scaled linearly with cell volume within the size range typical for an *E. coli* cell (~2μm, Sup. Fig. 1); therefore, this measure was used as a proxy for cellular size (**S**).

At the time of library construction ^11^, some bias was introduced by the investigators while arraying strains in plates. This bias resulted in the occasional presence of genes with similar functions together in the same plate, such as genes encoding chemotactic and flagellar proteins in plate #45 (Sup. Fig. 2). This bias, however, did not appear to significantly affect the mean fluorescence of the plate in relation to the dataset distribution (Sup. Fig. 3-6). Therefore, no further data normalization was performed. However, in light of this bias, the per-plate distribution of strains with significant phenotypes was further assessed (see below).

The average fluorescence of mVenus and mCherry varied approximately four-fold and was strongly correlated across the 3823 transformed KEIO strains (r = 0.90, Fig. 1A) and with the FSC measure (correlation 0.67 and 0.61 for mVenus and mCherry, respectively) (Fig. 1A). A change in cell size may globally affect cell physiology and indirectly affect heterologous gene expression in cases in which proteins are not sufficiently divided between daughter cells. This scenario was supported by the observed positive correlation between FSC and fluorescence output (Fig. 1A). We sought to assess the effects of gene knockout on heterologous gene expression after normalizing for the influence of variations in cell size (FSC). Fluorescence measurements were regressed against FSC to obtain the residuals for mCherry and mVenus, and the average of the two values was used as the FSC-normalized measure of heterologous gene expression **E**. To obtain a measure of the differential effect of KEIO knockouts on the individual reporter genes, the residuals of the FSC-regressed fluorescence using **E** as predictor were quantified. The resulting set of regressed mCherry and mVenus expression values were highly correlated with **E** (cor=0.94-0.98) and used as proxy for gene-specific effects (**G_spec_**).

**Figure 1.**
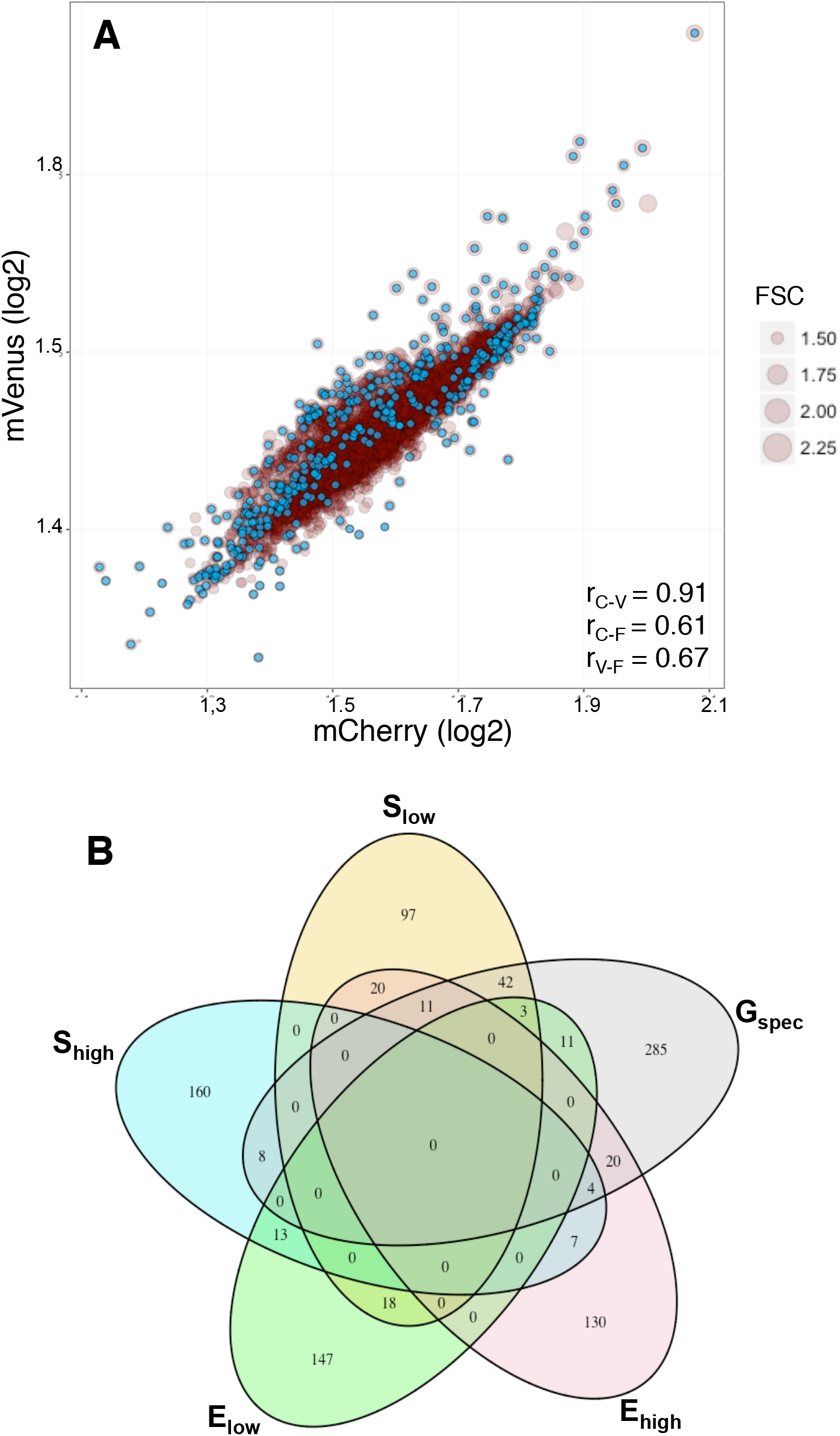
**(A)** Scatterplot of the mean population single-cell measure of cell volume (**FSC**) and fluorescence of mVenus and mCherry reporters (cyan dots: significant knockouts after FSC regression; r: Pearson correlation coefficients). **(B)** Venn diagram of the overlap between **S_high/low_**, **E_high/low_** and **G_spec_** extreme value genes.

### E and G_spec_ features are not affected by the same cell functions

The top and bottom 5% quantiles in each distribution were used to select genes with significant feature values. These extreme values were homogenously distributed across the dataset (Fig. 1A, cyan dots), thus indicating that the regression did not introduce noticeable bias. The number of genes in each group with higher or lower **S** or **E** values denoting **S_high_** /**S_low_** and **E_high_**/**E_low_** phenotypes was 153 and the number of genes with a G_spec_ phenotype was 384. Some co-localization of genes in KEIO plates was observed: genes with a **S_high_** phenotype were concentrated in the range between plate #17 and #43 (Sup. Fig. 7), several **E_high_** genes were found in plates #33, #37 and #39, and **E_low_** genes were found in plate #45 (Sup. Fig. 8). In addition, several genes with the G_spec_ phenotype were found in plates #61 and #89 (Sup. Fig. 9). There did not appear to be a clear link between the co-localization of genes in a certain plate and significant shifts in multiple features. For instance, although a number of genes with the **E_high_** - **S_high_** phenotype were found in the same #37 and #39 plates (Sup. Fig. 7-8), other plates, such as plate #33, contained a number of genes with **E_high_** but insignificant **S** or **G** phenotypes.

In general, a large number of genes had a significant value only in one feature. A **S_high_** phenotype combined with an **E_high/low_** shift was found only in ~7-8% of the genes, while a **S_low_** phenotype shared ~13% of the genes with the **E_low_** phenotype and a higher (~20%) fraction with the **E_high_** phenotype (p < 10^−4^, Fig. 1B). **E** and **G_spec_** phenotypes also shared 14-19% of genes. Notably, 33% (53/159) of genes with a **S_low_** shift also had a **G_spec_** phenotype compared with only 7.5% of those with a **S_high_** phenotype. This result may indicate that a decrease in cell size may disrupt the balance in the expression of two synthetic genes more significantly than an increase in cell size (Fig. 1B).

Next, the presence of functional enrichments in gene groups with specific **E** or **G** phenotypes was assessed. The **E_low_** group was enriched in knockouts in Enterobacterial Common Antigen (ECA) biosynthesis and flagella assembly (Bonferroni corrected p <10^−2^, Table 1 - bottom). We also found that knockouts for other membrane-associated functions, such as chemotaxis, cell adhesion, membrane transport and Two-Component Signal (TCS) transduction (KEGG pathways, p < 10^−1^), were enriched in this group. Genes with the **E_high_** phenotype were characterized by an enrichment in nucleotide biosynthesis activity for both purine and pyrimidine nucleotides (Table 1 - top). In this group, enrichment for homologous recombination functions, including the key genes *ruvA, priC* and *holD,* were also found. Purine nucleotide biosynthesis was also enriched among genes with **E_high_** − **G_spec_** phenotypes (p < 0.5) (Table 1). In this group, there were also KEIO strains of important housekeeping genes, such as *priB* and *priC,* which are involved in DNA replication. In addition, genes with key functions in mRNA (*pnp*) and protein stability (*cpxA*) as well as protein folding and export (*tatBC, dnaK, dmsD*) were identified (Table 1). Importantly, *tatBC* and *dmsD* physically interact in the cell and share a **E_high_** − **G_spec_** phenotype, but these KEIO knockouts were located in different KEIO plates (#89, #71 and #61) indicating that significant functional gene enrichment is not plate-biased.

**Table 1.**
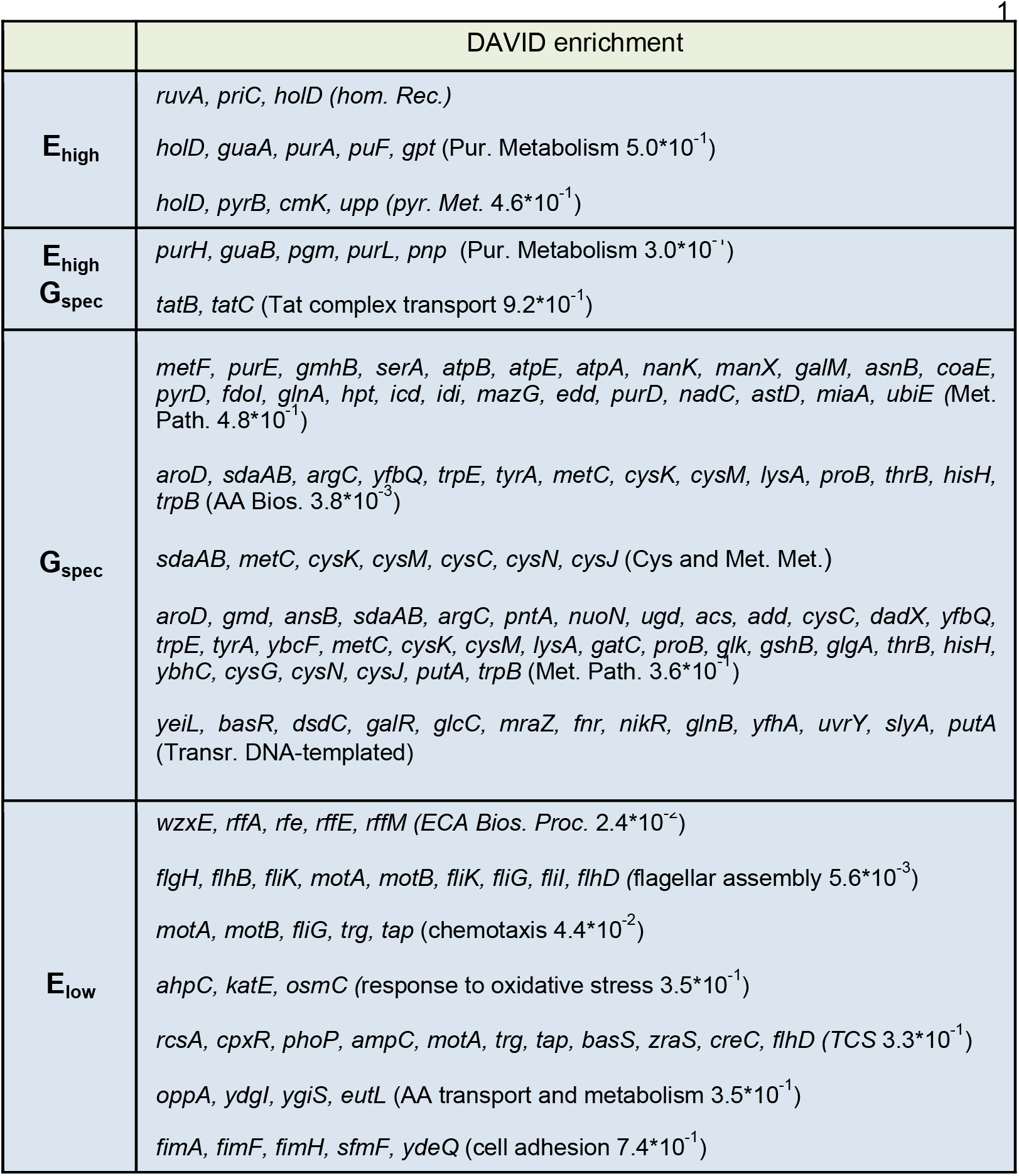
Significantly enriched GO Biological classes or KEGG pathways (Bonferroni-corrected p-value) within groups of genes with a significant shift in the **E** or **G_spec_** values.

Genes with a **G_spec_** phenotype belonged to a diverse range of cell functions including transcription factors and enzymes involved in central carbon metabolism and amino acid biosynthesis (p < 10^−2^). Overall, E and **G_spec_** phenotypes did not share substantial sets of cellular functions beyond genes involved in purine nucleotide biosynthesis.

### Cell function disruption triggers specific changes in S and E features

To gain a better understanding of the role of different cell functions on S and E, KEIO knockouts populating all observed combinations of **S_high/low_** and **E_high/low_** patterns were investigated. In addition, the presence of a **G_spec_** effect, which quantifies expression imbalance between the two reporter genes, was assessed in each of the observed **S**-**E** patterns. The final matrix consisted of 17 combinations populated by 400 genes with significant feature values (Fig. 2).

**Figure 2.**
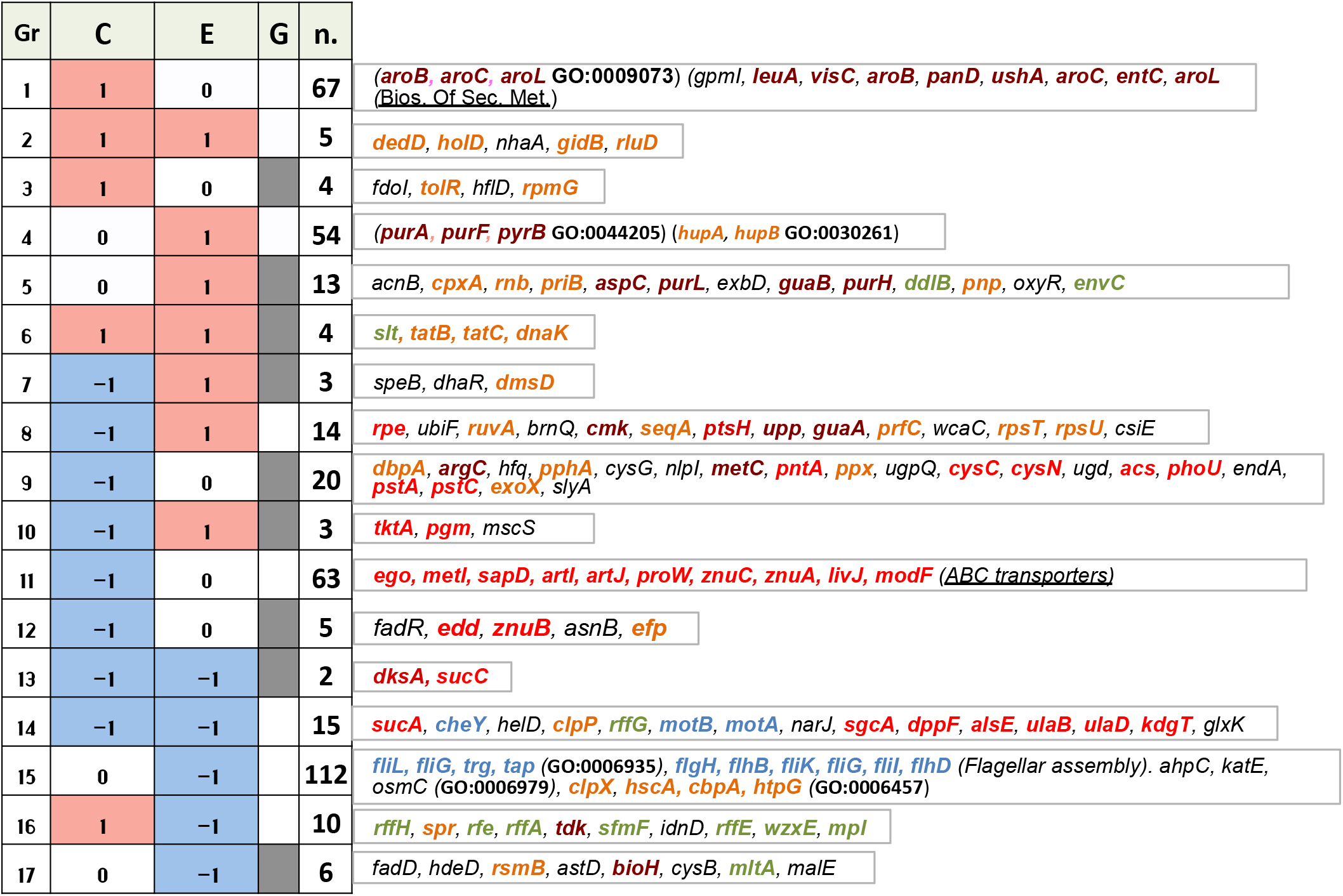
Distribution of genes in 17 phenotypic patterns (groups, **Gr**) with significantly (>2 st.dev.) increased (+1 /pink) or decreased (−1/cyan) cell size **(S)**, generic synthetic gene expression **(E)** and gene-specific effects **(G)**. For each combination of **S, G** and **E** patterns listed are either the individual extreme genes (for <20 genes), or the functional enrichments (DAVID) of Gene Ontology classes (bold/GO) or KEGG pathways (underscored). (Text color: brown = AA biosynthesis; orange = important to cell growth; red = nutrient uptake and catabolic reactions; cyan = motility and chemotaxis function; dark green = cell membrane structural component).

In general, the majority of genes (344 or 82%) showed a single-feature phenotype rather than significant changes in both features. Because there is a degree of co-localization of knockouts of genes with similar functions within the KEIO collection (see above), the plate aggregation of sets of genes sharing a specific **S-E** phenotype was investigated. Generally, strains from the entire KEIO collection populated each pattern (Sup. Fig. 10A-C and info). However, we found some co-localization of knockouts of genes involved in chemotaxis and flagellum biosynthesis in addition to a set of 4 protein chaperones (Group 15) on plate #45 (Sup. Fig. 10D). These genes shared a **E_low_** phenotype together with 7 other chemotactic/motility genes and the proteases ClpP/X, which resided in different KEIO plates. In conclusion, although we could not completely rule out a plate-effect for these functions, their influence on the Size-Expression interaction was supported by a larger number of members from the entire dataset.

The features-matrix was re-arrayed to observe the gradient of the shift with **S_high_** and **E_low_** placed at the top, bottom and all other possible combinations in between. This arrangement displayed a pattern that suggested that most genes were characterized by a similar up/down shift in the **S** and **E** features. Only 30 genes presented a mixed phenotype (for example **S_down_ - E_high_**), representing a total of only 7.5% of all cases. This result indicated a lack of substantial protein ‘concentration’ effect, likely a result of the regression against the FSC measure.

Next, all phenotypes with >20 members were assessed for functional enrichment, while gene lists were reported for smaller groups. Although a large number of genes presenting only a single-feature phenotype were observed earlier (Fig. 1B), the effect of disrupting major cell functions generally affected both **S** and **E** phenotypes simultaneously. Only the impairment in amino acid biosynthesis, and particularly in aromatic amino acids, appeared to specifically lead to a **S_high_** phenotype (Bonferroni corrected p < 10^−1^) (Fig. 2 group 1 brown text). Many genes with key housekeeping cellular functions presented a significant **S_high_** or **E_high_** phenotype either alone or in combination (Fig. 2, groups 2-6 orange text, see also Table 1). Intriguingly, as discussed above (Table 1), the effect of impairing nucleotide biosynthesis appeared to primarily lead to the E_high_ phenotype, with or without a **S_low_** or **G_spec_** effect (Fig. 2, groups 4, 5, 8 brown text). The observation that the deletion of amino acids and nucleotide biosynthetic enzymes showed an exclusively defined effect of increasing cell size or generic synthetic gene expression, respectively, is noteworthy and further discussed below.

Disruption in carbon metabolism and protein synthesis/folding appeared to dominantly be associated with **S_low_** and/or **E_low_** phenotypes. Knockout strains sharing a **S_low_** phenotype were involved in nutrient and metal ion uptake, including phosphate (*pstA, pstC*), sulfur (*cysC, cysN*) and zinc (*ZnuA, ZnuB*) (Fig. 2 - groups 8-10 red text). The **S_low_-E_low_** pattern was enriched with genes associated with carbohydrate catabolism (*sucA, sucC*) and a critical regulator of stationary phase onset (*dksA*). The lack of several cellular chaperones (Gene Ontology class GO:0006457, Bonferroni corrected p < 10^−1^) presented an exclusive **E_low_** phenotype, thus indicating that a lack of cellular protein folding functions negatively affected heterologous gene expression (Fig. 2 group 14, discussed below). Notably, these knockouts did not present a **G_spec_** effect; therefore, the effect on gene expression appeared to be global and did not affect individual genes differentially.

A **S_low_** - **E_low_** phenotype was also found for knockout strains for various genes involved in bacterial chemotaxis (*cheY, motA, motB*). However, disruptions in flagellum assembly (GO: 0006935 and KEGG pathway flagellar biosynthesis) were linked to an exclusive **E_low_** phenotype (Group 14, Fig 2, Bonferroni corrected p < 10^−1^). These findings indicated that cell motility and global gene expression are co-regulated in *E. coli.* Indeed, there is compelling evidence that widespread regulatory protein acetylation links nutrient scavenging and cell motility ^15^.

A disruption of ECA synthesis also showed an **E_low_** phenotype, but, in contrast to chemotaxis genes, this effect was combined and resulted in a **S_high_** shift (Fig. 2 - Group 16). Because generic cell growth delay appeared to lead to a **S_high_** − **E_high_** shift (Groups 2-3), the effects of deletions in ECA structural components were likely to have occurred through a different mechanism (discussed below) that led to defined and unique changes in the **S-E** features.

### A map of the Size-Expression interaction

Most cell functions played a role in the Size-Expression interaction defined by the **S** and **E** features. In this interaction, both features largely changed correspondingly (Fig 3, color gradient). Several KEIO knockouts with a **S_high_** - **E_high_** phenotype carried a disruption in key bacterial growth functions, including cell division, protein folding and DNA replication (Fig. 3 top-middle). However, a **S_high_** - **E_high_** phenotype was not common among all 32 knockouts of genes involved in cell division (GO:0051301), but among those responsible for outer membrane invagination (cytokinesis), in which the Tol-Pal system plays an important role ^16^ (Sup. Fig. 11). Bacterial DNA replication relies on a complex series of enzymatic and regulatory mechanisms. A significant number of KEIO strains of genes associated with the *E. coli* GO class ‘DNA-dependent DNA replication’ (GO:0006261) were found to be associated with a **S_high_** − **E_high_** phenotype (p < 10^−4^, 5 of 9 Sup. Fig. 12). The relationship between cytokinesis and chromosome replication is not completely clear, but it is reasonable to conjecture that growth delay (**S_high_**) may indirectly increase cellular fluorescence (**E_high_**) in these mutants. However, knocking out *seqA,* a negative modulator of chromosome replication, led to a **S_low_** - **E_high_** phenotype, thus suggesting a more direct mechanism.

**Figure 3.**
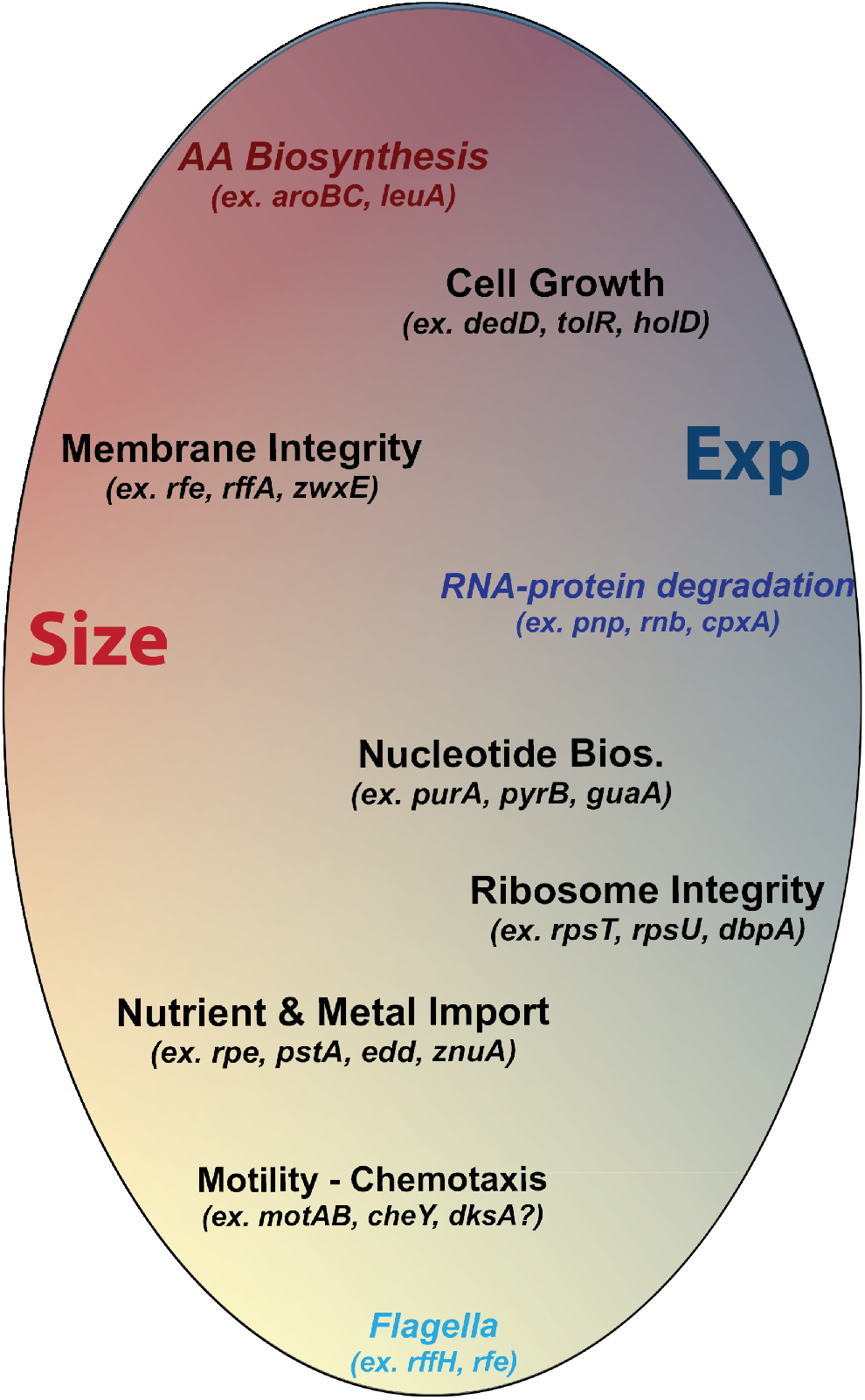
A map of the effect of major cellular systems on the interaction between cell **Size** (red) and synthetic gene **Exp**ression (blue). The impact of each cell functional category can be to increase (darker shade) or decrease (lighter shade) predominantly one side of the interaction or both (represented via the positioning towards respectively the edge or the inner of the oval).

For a number of strains, cell size and global gene expression did not vary together. For example, some KEIO knockouts with a **S_low_** - **E_high_** phenotype indicated an important role in the cellular SOS response (Fig. 2 Groups 8-9). However, a lack of other members associated with this DNA repair function did not result in a univocal **S-E** phenotype (p = 0.17, Sup. Fig. 13). Other knockouts with a **S_low_** - **E_high_** pattern did not have ribosome structural components (*rpsT, rpsU, dbpA*) or important steps of RNA translation (*prfC, efp*) (Fig 3 - mid right). Twelve knockouts in ribosome structural genes were present in the dataset and, significantly, 5 of them - *rpsU, rpsT, rpmJ, rpmE, rplA* - showed a **S_low_** − **E_high_** phenotype (p = 0.02, Sup. Info.). After the analysis was extended to all genes important for cellular translation (GO:0006412, 26 genes in dataset), the link to a **S_low_** − **E_high_** pattern was even more substantial (p < 10^−2^) (Sup. Fig. 14). Importantly, 18/26 of these genes also showed a significant **G_spec_** phenotype (p < 10^−4^), thus suggesting that major gene expression disruption also differentially affects the expression of heterologous genes apart from causing a small-cell phenotype.

The second important aspect of the Size-Expression interaction was that most cell functions affected both sides. Indeed, only a limited set of related KEIO gene knockouts showed an exclusive **S** or **E** phenotype. For example, a number of structural flagellar components and various cellular chaperone proteins showed a predominantly **E_low_** phenotype (Fig 3 - cyan). An extension to all members of the Gene Ontology term class ‘chemotaxis’ (GO:0006935, 23 genes) and ‘bacterial-type flagellum’ (GO:0009288, 23 genes) also yielded a significant decrease in E (p < 10^−4^ and p < 0.005, respectively, calculated by bootstrapping) (Sup. Fig. 15). Only a few knockouts in this group (e.g., *flhE, flgG, fliQ,* Sup. Fig. 15) had an associated **S_high_** value, a pattern reminiscent of that of knockouts of genes involved in ECA biosynthesis (Fig 2 Group 16) (Fig. 3 middle left). The inclusion of the 13 other gene members of the ‘Enterobacterial Common Antigen Biosynthetic Process’ in the dataset further supported the phenotype for this group of genes (GO:0009246) (p < 10^−4^, Sup. Fig. 15).

In contrast, KEIO knockouts probably leading to increased RNA stability, such as those in ribonuclease II (*rnb*) and polynucleotide phosphorylase (PNPase), which degrade various types of mRNA ^17^, presented an exclusive **E_high_** phenotype (Fig. 3 - dark blue). Finally, the lack of a number of amino acid biosynthetic genes, and particularly those for aromatic amino acids, presented a unique **S_high_** phenotype (Fig. 3 top), which was confirmed after the analysis was extended to all genes associated with the ‘Cellular amino acid biosynthetic process’ GO class (GO:0008652, p = 0.011 calculated by bootstrapping, 86 genes in the dataset) (Fig. 3 - dark red).

## Discussion

The present study describes how specific cellular activities influence the relationship between a cell’s ability to grow and its synthesis of neutral heterologous proteins. We found that bacterial cell functions could be broadly subdivided into 4 systems of clearly defined phenotypes: Membrane system, Ribosome-RNA-Protein stability (RRP) system, the Biosynthetic system and the Energy-Motility system. Mutations in the Membrane system mainly lead to an increase in cell size and are likely to have indirect effects on protein concentration, which manifest as changes in gene expression. The RRP stability system primarily affects gene expression, with or without a global (size) effect that depends on the structural nature (e.g., ribosome) of the disruption. The lack of certain steps in the Biosynthetic system, particularly those for amino acids, are likely to have a global impact simulating the bacterial stringent response with a broad remodeling of cellular protein composition ^18^. Finally, disruptions in the Energy-Motility system, which is highly interconnected in the stationary phase by the global regulators ppGpp and dksA ^19^, may exert a negative feedback on the ability of the cell to search for nutrients or increase gene expression, for example via global protein acetylation ^20,21^, which leads to a small-cell low-expression phenotype.

This study also shows that most mutations affecting cell functions, including cell motility, membrane structure, chromosomal DNA replication, repair and homologous recombination, do not differentiate between identically expressed synthetic genes (no **G_spec_**). Not surprisingly, however, gene knockouts with differential effects on the two reporter genes were primarily related to protein expression and were associated with functions including ribosome structure, translation (*efp*), protein folding (*cpxA, dnaK*) and transport (membrane TAT complex).

An important challenge in both metabolic engineering and synthetic biology is to precisely understand how the introduction of engineered or non-native components into a biochemical network influences the behavior of the entire system.

This study mapped how the removal of each cell function in *E. coli* influences the cell, the synthetic genes introduced, or both. This accurate systematic analysis of the cellular context of synthetic gene expression may facilitate metabolic engineering workflows and systems-level modeling of this model prokaryote that serves as a key industrial workhorse organism.

## Methods

### Strains, plasmids and media

Single-gene knockout strains were obtained from the KEIO collection and wild type laboratory E. coli from the Joint Bio-Energy Institute (JBEI, Emeryville-CA). For construction of plasmid library, strains were cultivated in LB media supplemented with appropriate antibiotics and subjected to chemical (CaCl_2_) transformation. For flow cytometry and further analysis, all strains were cultured in Neidhardt’s MOPS-based Rich defined medium (Teknova), supplemented with 0.5% glucose and antibiotics Ampicillin or Kanamycin (50-70 □g/ml). The construction of pEZ8-123 synthetic genetic probe has been previously described ^10^.

### Flow Cytometry

All data were acquired with a Guava Flow Cytometer (FC) as follows: 24-36 strain cultivations at a time (2-3 rows for each 96 well plate) were inoculated at a 1:80 dilution from overnight-cultures in warm MOPS rich medium supplemented with 0.5% glucose, and grown for exactly 1h15’, in a shaker incubator at 37C. At the end of the incubation, the OD of the culture was read with the help of a microtiter plate reader to determine the number of cultures at mid-exponential phase (a 0.3-1.0 range corresponded to a spectrophotometer OD reading of ~1.3-2.0). These cultures were diluted 1:200 in PBS + translation inhibitor, and single-cell readings acquired via flow cytometry.

### Statistical and computational analysis

Programming software R was utilized for statistical and other type of data analysis. Please refer to supporting information for a detailed description of methodologies and functions.

## Supporting information

Refer to the web version for supplementary material and information. This file provides in depth description of experimental and statistical methodology, as well as supporting figures and tables.

## Acknowledgments

I would like to thank Prof. Adam Arkin (University of California-Berkeley, USA) for financial and intellectual support throughout the study. Without inspirational and extensive discussions with Prof. Arkin over the years this study could not have been completed. I also want to thank Dr. Marcin Joachimiak for protracted dialogues on the appropriate analysis and results significance.

This work was funded by the National Science Foundation as part of the Synthetic Biology Engineering Research Center grant number 04570/0540879.

